# Ornaments for Accurate and Efficient Allele-Specific Expression Estimation with Bias Correction

**DOI:** 10.1101/2023.10.25.564046

**Authors:** Abhinav Adduri, Seyoung Kim

## Abstract

Allele-specific expression has been used to elucidate various biological mechanisms, such as genomic imprinting and gene expression variation caused by genetic changes in *cis*-regulatory elements. However, existing methods for obtaining allele-specific expression from RNA-seq reads do not adequately and efficiently remove various biases, such as reference bias, where reads containing the alternative allele do not map to the reference transcriptome, or ambiguous mapping bias, where reads containing the reference allele map differently from reads containing the alternative allele. We present Ornaments, a computational tool for rapid and accurate estimation of allele-specific expression at unphased heterozygous loci from RNA-seq reads while correcting for allele-specific read mapping bias. Ornaments removes reference bias by accounting for personalized transcriptome, and ambiguous mapping bias by probabilistically assigning reads to multiple transcripts and variant loci they map to. Ornaments is a lightweight extension of kallisto, a popular tool for fast RNA-seq quantification, that improves the efficiency and accuracy of WASP, a popular tool for bias correction in allele-specific read mapping. Our experiments on simulated and human lymphoblastoid cell-line RNA-seq reads with the genomes of the 1000 Genomes Project show that Ornaments is as efficient as kallisto, an order of magnitude faster than WASP, and more accurate than WASP and kallisto. In addition, Ornaments detected genes that are imprinted at transcript level with higher sensitivity, compared to WASP that detected the imprinted signals only at gene level.

## Introduction

Allele-specific expression has been used to characterize various biological phenomena in diploid organisms, including gene expression affected by *cis*-acting variants in an allele-specific manner ^1,2^, allele-specific nonsense-mediated mRNA decay ^3,4^, and monoallelic expression of imprinted genes ^5,6^. Allele-specific expression is typically measured as RNA-seq read depths at heterozygous loci. There are two well-known biases introduced in allele-specific read mapping: reference bias arising from reads with alternative alleles that do not map to the reference transcriptome; and ambiguous mapping bias arising from reads that map to a heterozygous site but also to homozygous sites in other genomic locations, causing the same reads with different alleles to map to different genomic locations ^7,8,9^.

Removing these biases efficiently for accurate allele-specific expression estimation has been a challenging problem. Mapping reads to a diploid personalized transcriptome ^10,11^ removed only reference bias but not ambiguous mapping bias. WASP ^7^, a popular tool for removing ambiguous mapping bias, was not adequate, as it simply discarded ambiguously mapped allele-specific reads, obtained allele-specific read counts only at gene level but not at transcript level, and was computationally costly for both the initial read mapping to the reference genome and the re-mapping for bias correction. Kallisto ^12^, a popular tool for rapid transcriptome quantification, probabilistically assigned multi-mapped reads given a diploid transcriptome with known variant phases. However, with kallisto and other related methods ^13,14^, inaccurate phasing could lead to inaccurate allele-specific signals.

Here, we introduce Ornaments, a tool for accurate and rapid estimation of allele-specific transcript expression at unphased heterozygous loci from RNA-seq reads. Ornaments removes reference bias by accounting for sample-specific variant information, and ambiguous mapping bias by probabilistically assigning reads to multiple transcripts and variant loci they map to.

Ornaments is a lightweight modification of the two stages of kallisto ^12^, the read mapping and quantification stages, that improves the accuracy and efficiency of WASP as well as the accuracy of kallisto, while maintaining the computational efficiency of kallisto. In the read mapping stage, Ornaments uses an ornament transcriptome de Bruijn graph (tDBG), a new data structure obtained by augmenting the colored tDBG of kallisto with ornaments for variants (Figs. 1a and 1b). Each ornament has two shades representing the two variant alleles. With this ornament tDBG, Ornaments performs variant-aware pseudoalignment of each read, a modification of kallisto pseudoalignment, to find a variant-aware equivalence class defined as the set of transcripts and variant alleles that the read maps to (Figs. 1c and 1d). In the quantification stage, Ornaments uses a modification of the kallisto mixture model to obtain expected allele-specific read counts at heterozygous loci in addition to transcript quantification. In our experiments on simulated and human lymphoblastoid cell-line RNA-seq reads ^15^ with the genetic variants of the 1000 Genomes Project samples ^16^, Ornaments was nearly as efficient as kallisto and was an order of magnitude or on average 26 times faster than WASP. In addition, Ornaments improved the accuracy of WASP by adequately removing read mapping bias at transcript level and the accuracy of kallisto by accounting for genetic variants.

**Figure 1:**
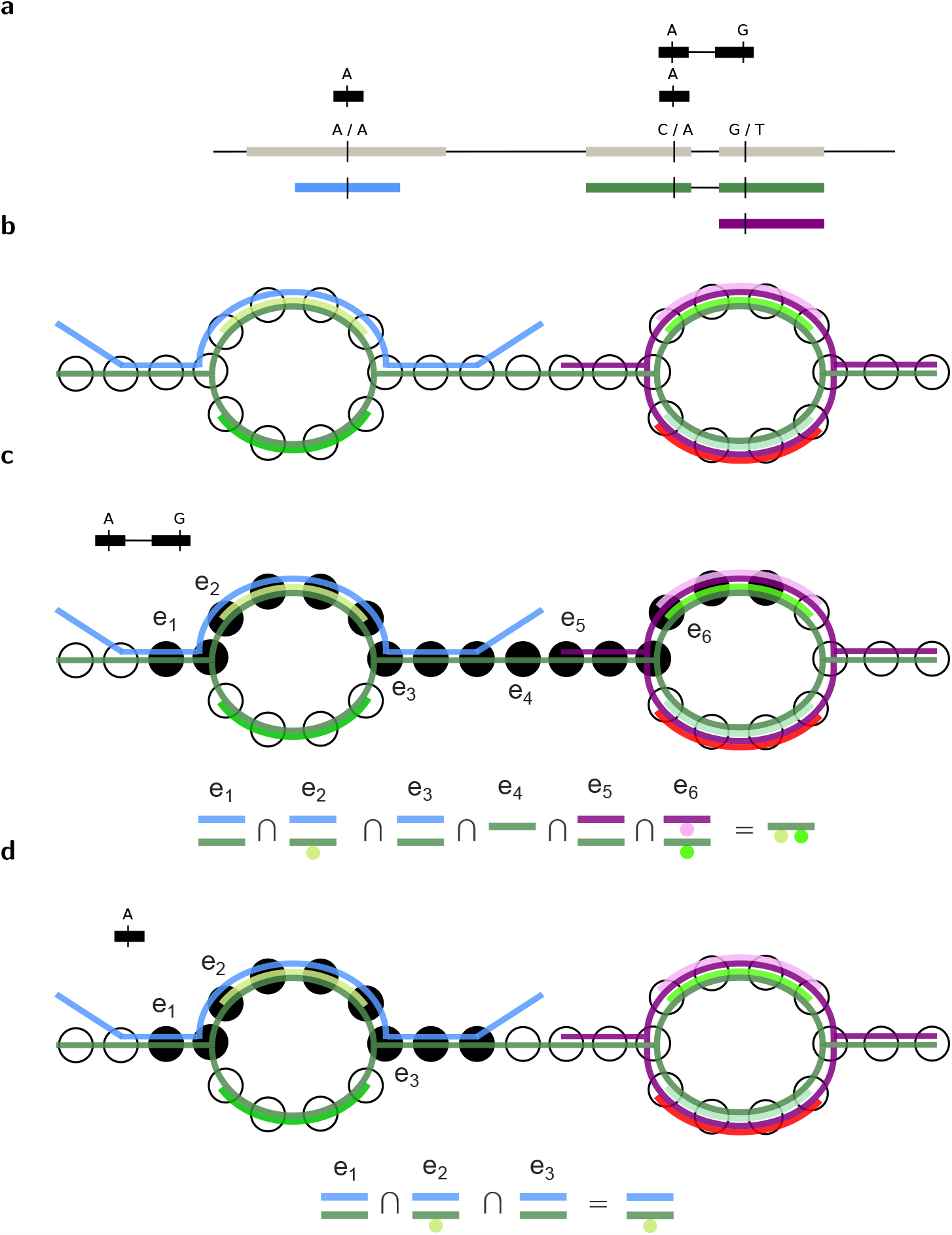
Ornaments overview. (a) The genome (grey) with three transcripts (blue, green, and purple), two heterozygous SNPs, one uniquely-mapped allele-specific read (top read) and one ambiguously-mapped allele-specific read (bottom read mapped to two repeat regions in blue and green transcripts). (b) An ornament tDBG consists of a colored tDBG, where *k*-mer nodes are colored for transcripts, and ornaments, where *k*-mer nodes are shaded for variant alleles. (c) The variant-aware pseudoalignment of the uniquely-mapped read finds the variant-aware equivalence class for the read as the intersection of the sets of colors for transcript contigs, and the union of shades associated with the colors in the intersection. The first node of each contig overlapping with the read is marked as *e*_1_, …, *e*_6_. (d) The variant-aware pseudoalignment of the ambiguously-mapped read.

## Material and methods

We introduce Ornaments and describe the two stages of the Ornaments method, the first stage for variant-aware pseudoalignment of reads with an ornament tDBG and the second stage for allele-specific expression estimation with a mixture model. We describe how to prepare variant information to construct an ornament personalized transcriptome that is used as an input to Ornaments.

### Preparation of variant information

We prepare variant information as follows. Given the reference transcriptome, transcript annotation, and information on the genomic coordinates and alleles of variants, we extract SNPs and indels in exonic regions, and transform the genomic coordinates of those variants into transcriptomic coordinates. Then, for each variant, we store the information on the transcript it appears in, the transcriptomic coordinates of the variant, and the allele. When a variant appears in multiple alternatively spliced transcripts, a single variant with a single genomic coordinate is associated with multiple transcriptomic coordinates. If there are multiple overlapping indels at the same genomic coordinate, we only keep the last one.

### Ornament personalized transcriptome

From the variant information above and reference transcriptome, we prepare an ornament personalized transcriptome, which will be used to build an ornament tDBG over *k*-mers for an Ornament index. An ornament personalized transcriptome consists of transcript sequences and ornament sequences. The transcript sequences are set to those of the reference transcriptome with a modification to alternative allele at each alternative-allele homozygous locus, and retain their transcript names in the reference transcriptome. Two ornament sequences are added for each pair of a heterozygous variant and transcript containing this variant, where each ornament sequence consists of each variant allele and the flanking sequences of length *k* on each side of the variant in the transcript. In rare cases of *n* heterozygous loci within *k* base pairs, where almost always *n*=2, an ornament sequence is added for each of the 2^*n*^ combinations of the alleles. For multiple overlapping indels, an ornament sequence is added only for the first indel. The ornament sequences are named as a concatenation of the name of the transcript of origin, the position of the variant within the transcript, and the allele.

The construction of ornament personalized transcriptome is efficient in both space and time. It takes a few seconds to construct it, and its size is not significantly larger than the size of the reference transcriptome, since ornament sequences are typically substantially shorter than transcript sequences.

### Constructing ornament index

With an ornament personalized transcriptome as input, we construct an ornament index that consists of an ornament tDBG and ornament hash table. An ornament index is implemented as an outer layer over kallisto index. To construct an ornament tDBG (Figs. 1(a) and 1(b)), we apply the kallisto routine of constructing a colored tDBG to both transcript and ornament sequences in the ornament personalized transcriptome. Colors are assigned to both transcript and ornament sequences and each *k*-mer node in the tDBG is annotated with the set of colors corresponding to the transcript or ornament sequences it appears in. We call it an ornament tDBG because the colors for the *k*-mers of ornament sequences appear as ornaments in the colored tDBG. The colors of the ornament sequences are mapped to shades using an ornament hash table as we describe below.

We construct an ornament hash table that maps a *k*-mer to a set of transcripts and variant alleles. An ornament hash table consists of a kallisto hash table, which maps each *k*-mer to a set of colors corresponding to contigs of transcripts the *k*-mer appears in, and an auxiliary hash table, which maps the color to an ornament shade if the color corresponds to an ornament sequence. The auxiliary hash table stores two pieces of information for each ornament shade: the location of the variant within the transcript and the variant allele at that locus.

### Variant-aware pseudoalignment

Using the ornament index, we map RNA-seq reads to personalized transcriptome. We modify the pseudoalignment of kallisto to variant-aware pseudoalignment, which is an assignment of a read to a variant-aware equivalence class (Figs. 1(c) and 1(d)). A variant-aware equivalence class for a read is defined as the set of possible transcripts and alleles of origin for the given read and is obtained in two steps. In the first step, each *k*-mer of the read is mapped to a set of colors via the kallisto hash table, and if the *k*-mer maps to an ornament sequence, is also mapped to ornament shades via the auxiliary hash table. Then, in the second step, we combine the colors and shades of all *k*-mers of the read from the first step as follows. We first obtain a pseudoalignment of the read to a set of transcripts by finding the intersection of the sets of colors for all *k*-mers as in kallisto. Then, we obtain a variant-aware pseudoalignment by taking the union of the shades for the transcript colors included in this intersection. A shade is included in the variant-aware pseudoalignment of a read only if the read contains all *k*-mers of the given shade in the ornament tDBG, unless the *k*-mers mapped to the shade are located in either end of the read.

Applying variant-aware pseudoalignment to all reads from a sample results in read counts for each of the variant-aware equivalence classes. These are the sufficient statistics needed for quantification of transcript expression and allele-specific expression at heterozygous loci.

### Quantification of transcript expression and allele-specific expression at heterozygous loci

Given these read counts for variant-aware equivalence classes, we extend the mixture model and EM algorithm of kallisto to quantify allele-specific expression at each heterozygous locus in addition to transcript expression. We provide a statistically rigorous set-up of the model and derivation of the EM algorithm that differ from those in kallisto, and show that our approach leads to the same transcript expression quantification as kallisto.

Let *E* ∈ {1, …, *N*_*E*_}, where *N*_*E*_ is the number of variant-aware equivalence classes, denote a random variable for the variant-aware equivalence class of a read. Let *T* ∈ {1, …, *N*_*T*_}, where *N*_*T*_ is the number of transcripts, be a random variable for a transcript label. We set up a mixture model for the variant-aware equivalence class of a read

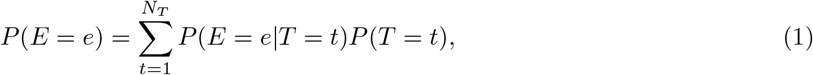

where the mixture component model *P* (*E* = *e*|*T* = *t*) and mixture proportion *P* (*T* = *t*) are given as

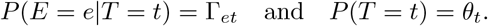

The mixture proportion parameter 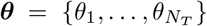 satisfies 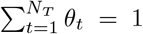, and models the unknown expression quantification for *N*_*T*_ transcripts to be estimated. In the mixture component model, Γ_*et*_ is the (*e, t*)th element of the *N*_*E*_ *× N*_*T*_ matrix **Γ** and is defined as follows:

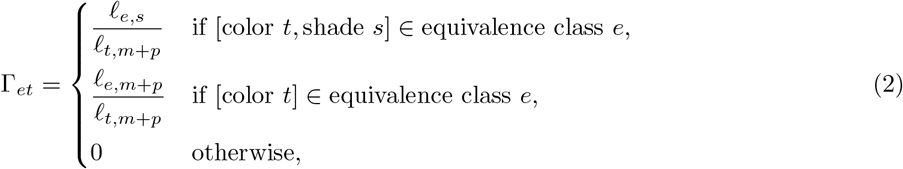

where *f*_*t,m*+*p*_ in the denominator is the combined length of the two alleles of the transcript *t, f*_*e,m*+*p*_ in the numerator is the combined length of the two alleles of transcript contig corresponding to the equivalence class *e*, and *f*_*e,s*_ in the numerator is the length of the contig corresponding to the equivalence class *e* and the variant allele *s*. Notice that each row of **Γ** has an identical numerator and each column of **Γ** has an identical denominator in all non-zero entries. It is straightforward to verify that the column sum of **Γ** is equal to 1, i.e., 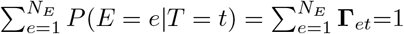.

Eq. (1) defines a proper generative model for the variant-aware equivalence classes of reads. To generate a variant-aware equivalence class for a read, transcript *t* is selected with probability *θ*_*t*_ and then a contig for equivalence class *e* in transcript *t* is selected with probability Γ_*et*_, which is proportional to the length of the contig as defined in Eq. (2). In contrast, the kallisto mixture model is defined with the contig length *l*_*e,m*+*p*_ = 1 in the numerator of Γ_*et*_, and thus, does not provide a proper description of the generative process. As we show below, it is not necessary to obtain the contig lengths *l*_*e,m*+*p*_ and *l*_*e,s*_ explicitly, since these quantities do not appear in the update equations of the EM algorithm. Thus, both our and kallisto’s model set-ups lead to identical parameter estimates.

We estimate the model parameters ***θ*** with the EM algorithm. For each read *r*_*i*_, where *i* = 1, …, *N*_*R*_ for *N*_*R*_ reads, let PA(*r*_*i*_) denote the variant-aware equivalence class of read *r*_*i*_. The log-likelihood of the data PA(*r*_*i*_)’s is

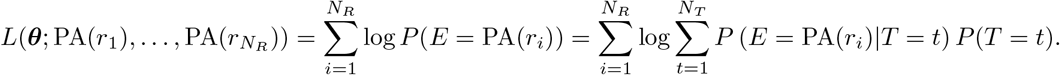

To estimate ***θ***, instead of directly maximizing the data log-likelihood above, the EM algorithm maximizes the expected complete-data log-likelihood:

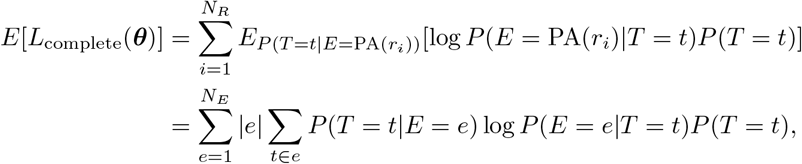

where |*e*| represents the number of reads assigned to the equivalence class *e*.

In each iteration of the EM algorithm, given the estimate 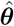 from the M step of the previous iteration, the E step computes

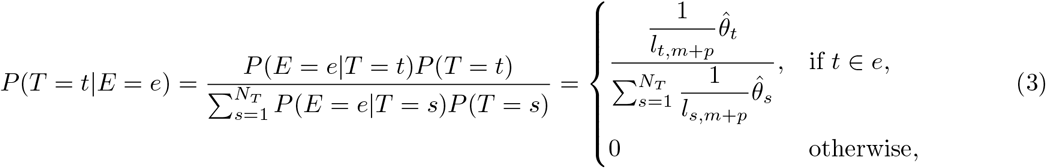

and given the above from the E step, the M step maximizes

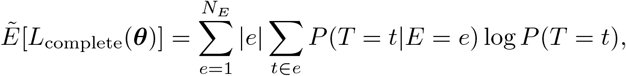

and updates the estimate as

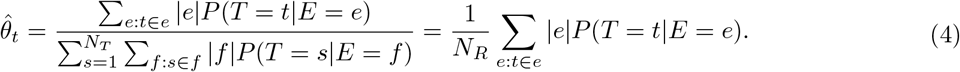

The second equality above holds from

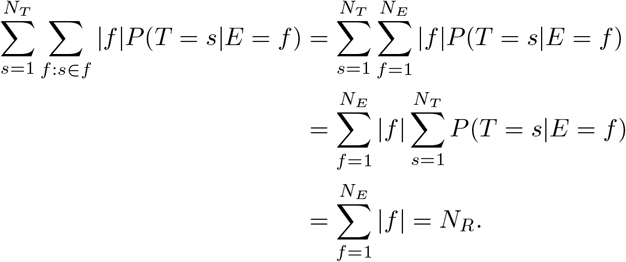

Notice that the contig lengths *ℓ*_*e,s*_ and *ℓ*_*e,m*+*p*_ of equivalence class *e* appear in neither the E-step update in Eq. (3) nor the M-step update in Eq. (4), and thus are not needed to estimate ***θ***. Convergence is called when the relative change for each *θ*_*t*_ is less than 0.1%.

Given the estimate 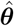, we quantify the allele-specific expression at each heterozygous variant locus as the expected allele-specific read depths. At heterozygous locus *j* for transcript *t*, we compute the expected read depths 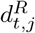 for the reference allele and 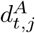 for the alternative allele by summing the expected read depth for each read in Eq. (3) over all reads:

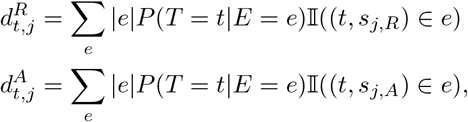

where 𝕀((*t, s*_*j,R*_) ∈ *e*)=1 if the pair of color *t* and shade *s*_*j,R*_ for the reference allele belongs to variant-aware equivalence class *e*, 𝕀((*t, s*_*j,R*_) ∈ *e*)=0 otherwise, 𝟙 and ((*t, s*_*j,A*_) ∈ *e*) is defined similarly for the alternative allele.

## Results

We benchmarked Ornaments against WASP and kallisto, using simulated and lymphoblastoid cell-line RNA-seq reads ^15^ for 165 individuals with SNP genotypes from the 1000 Genomes Project ^16^. These individuals were children of trios with known parental genotypes in the 1000 Genomes Project and thus with known haplotypes. We used the transcript annotation from GENCODE (version 36, Ensembl 102) to build ornament personalized transcriptomes. Low-quality RNA-seq reads that are too short or contain ambiguous nucleotides were removed using Trimmomatic 0.35^17^.

In all experiments, default settings were used for WASP and kallisto. In WASP, we used the STAR aligner ^18^ for the initial read mapping to the reference genome. We mapped reads to the transcriptomic regions of genomes by first mapping reads to the full genome with the WASP mappability pipeline and STAR aligner and then retaining only the reads that map to exonic regions or multi-map to at least one exonic region. To ensure that reads that map to SNP loci with alternative alleles are not dropped due to reference bias, when running the STAR aligner, we allowed reads to multi-map across up to 40 loci. Since the WASP re-mapping pipeline drops reads that are mapped to indels by the STAR aligner, in our comparison between WASP and Ornaments, we did not include reads that Ornaments map to indels and to SNPs within average read-length distance or 100bp of an indel.

### Simulation

For simulation study, we selected 10 samples among the 165 children and generated 60 million RNA-seq reads for each sample from the phased diploid transcriptome of the sample and ground-truth allele-specific transcript abundances. These 10 samples spanned multiple ethnicities (HG00405, HG00526, HG00709, HG00621 for East Asian ancestry; NA12766, NA12335, NA07029 for European ancestry; and NA18869, NA18930, NA19211 for African ancestry). The ground-truth allele-specific expression levels and background noise levels were set to the estimates obtained by applying RSEM ^19^ to the lymphoblastoid cellline reads that were aligned to the personalized transcriptome using Bowtie 2.0. The background noise was estimated to be 20% on average across samples. During simulation with the RSEM-simulate-reads program, we recorded reads overlapping with variants, from which we inferred the ground-truth allele-specific read counts at each heterozygous locus.

We first compared the number of reads dropped by different methods, as these reads can affect accuracy. WASP dropped on average three times as many reads per sample as Ornaments, and kallisto with reference transcriptome dropped 8.1% more reads than Ornaments (Fig. S1). Most of the reads dropped by kallisto and Ornaments were not allele-specific and were dropped because pseudoalignment requires exact *k*-mer matches. However, 95.5% of the reads dropped by WASP were ambiguously mapped allele-specific reads, and these dropped reads reduced the accuracy of WASP as we show below.

Ornaments outperformed WASP in the accuracy of allele-specific expression at heterozygous SNP loci, as it can correctly map and apportion ambiguously mapped allele-specific reads. The transcript-level estimates from Ornaments were summed over transcripts from the same gene, and these gene-level aggregates were compared to the corresponding gene-level read depths from WASP, at SNP loci with and without ambiguously mapped allele-specific reads. To determine if a SNP is involved in ambiguous read mapping, we used the variant-aware equivalence classes from Ornaments. If a variant-aware equivalence class that contains the shade/color pair for the given SNP also contains colors with no paired shade for other transcripts, then we determined that the SNP was involved in ambiguous read mapping. At SNP loci with ambiguously mapped allele-specific reads, Ornaments had significantly lower mean absolute relative difference for the estimated allele-specific expression (Fig. 2a) and higher correlation between the true and estimated allelic ratios than WASP (Fig. 2b). This in turn led to slightly higher accuracy for Ornaments at the other SNP loci without ambiguously-mapped reads (Figs. 2c and 2d).

**Figure 2:**
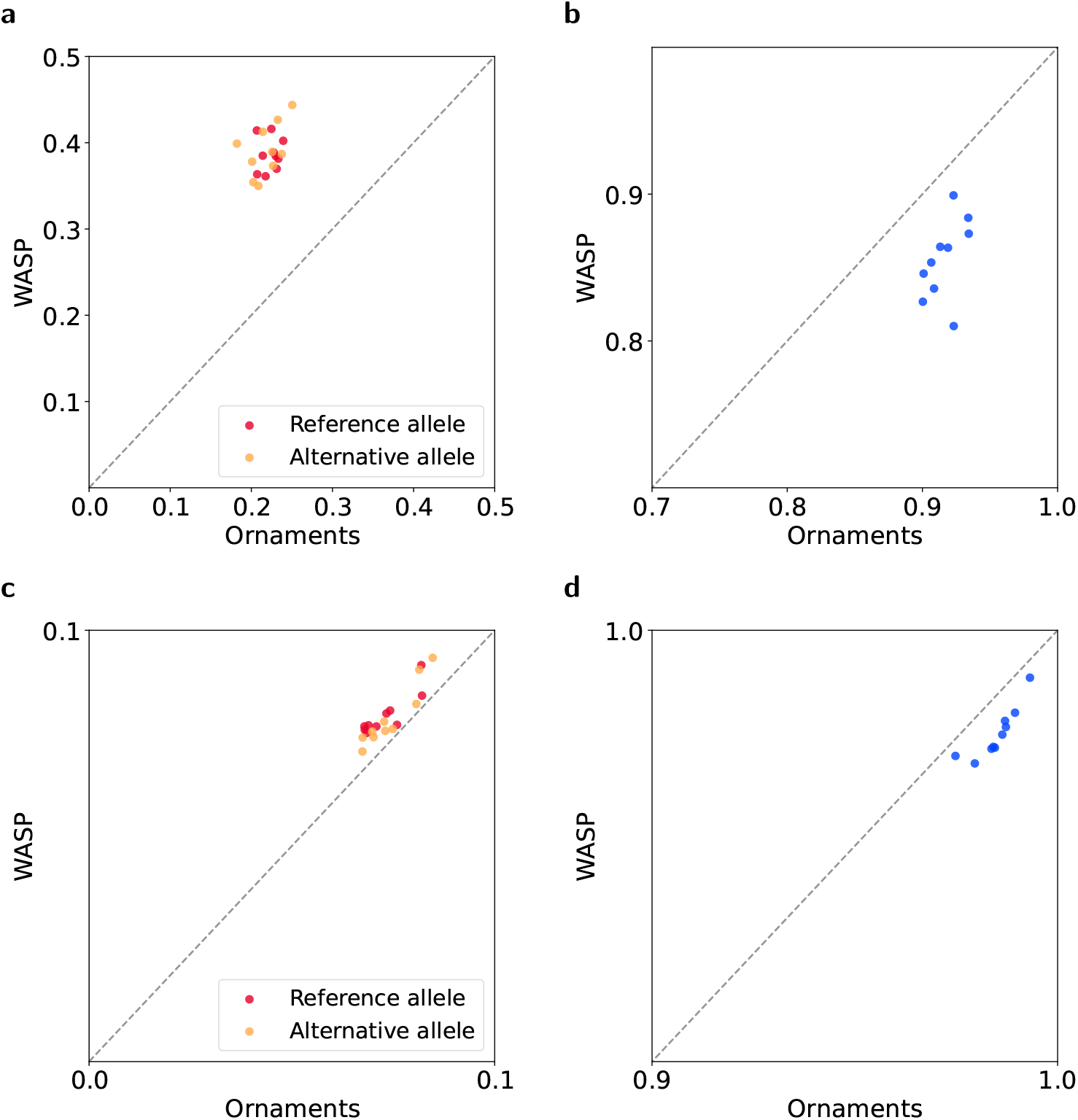
Comparison of Ornaments and WASP on the accuracy of allele-specific expression at heterozygous SNP loci using simulated data. At heterozygous SNP loci with ambiguously mapped allele-specific reads, (a) the accuracy of allele-specific expression, measured as mean absolute relative difference between the truth and estimated and (b) the accuracy of allelic ratios, measured as correlation between the true and estimated allelic ratios across loci. At heterozygous SNP loci without ambiguous allele-specific read mapping, (c) the accuracy of allele-specific expression and (d) the accuracy of allelic ratios. Each dot represents each of 10 samples.

Ornaments achieved higher accuracy than WASP in downstream analysis of detecting genes with differentially expressed alleles (Fig. 3). Given the allele-specific expression at heterozygous loci from Ornaments and WASP, we used the known variant phases to combine allele-specific signals across multiple loci within the same gene, and detected genes whose alleles are differentially expressed using GeneiASE ^20^ (*p*-val *<* 0.05). These genes were compared against the genes obtained by applying GeneiASE to the ground-truth allele-specific read counts. Across all 10 samples, Ornaments had higher sensitivity and specificity for detecting genes with differentially expressed alleles, suggesting that highly accurate allele-specific signals from Ornaments lead to higher accuracy in downstream analysis compared to WASP that discards ambiguously mapped allele-specific reads.

**Figure 3:**
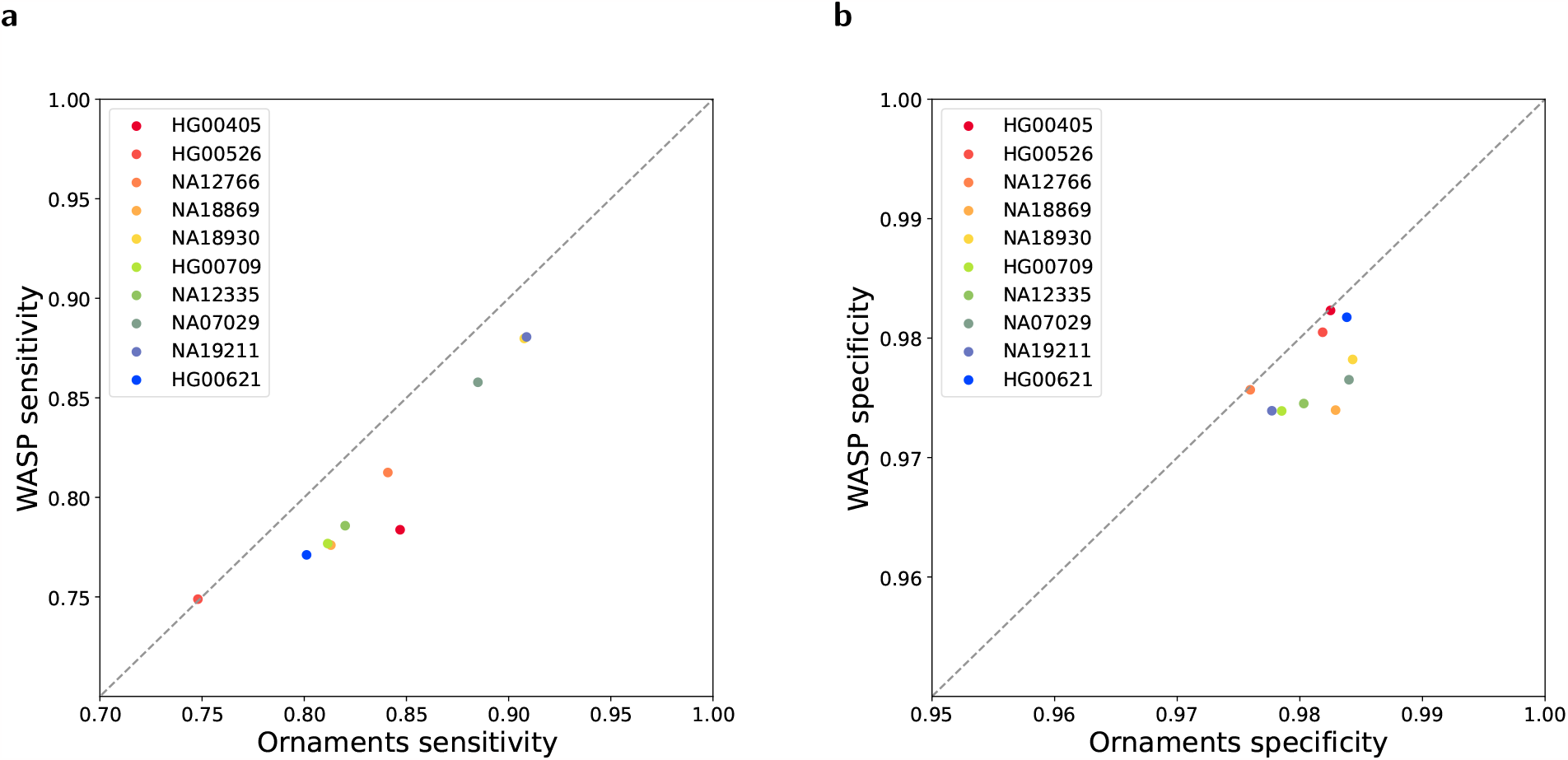
Comparison of Ornaments and WASP on the accuracy of detecting allele-specifically expressed genes using simulated data. Genes whose alleles are differentially expressed (*p*-value *<* 0.05) were obtained by applying GeneiASE to the allele-specific signals from Ornaments and WASP at each heterozygous SNP locus. These genes were compared against the genes obtained by applying GeneiASE to the ground-truth allele-specific read counts. (a) Sensitivity and (b) specificity of Ornaments and WASP.

Ornaments outperformed kallisto on the accuracy of transcript expression quantification, as kallisto cannot correctly map and apportion ambiguously mapped allele-specific reads. For 10 read datasets from repeated simulations from the genome of each sample, the accuracy was compared on expressed transcripts (true abundance *>* 0) using mean absolute relative difference, and on unexpressed transcripts using mean absolute difference, as mean absolute relative difference is known to be biased by small ground-truth values ^21^. Ornaments substantially outperformed kallisto when transcripts contained ambiguously mapped allele-specific reads (Figs. S2-S5), because kallisto without variant information cannot correctly map and apportion reads across the multiple mapped loci that involve both heterozygous and homozygous loci in repeat regions. Transcripts were considered as overlapping with ambiguously mapped reads, if any of the equivalence classes in Ornaments that contain the transcript also contains shade/color pairs in addition to colors alone. For transcripts without ambiguously mapped reads, Ornaments only slightly outperformed kallisto, even when the transcripts contained variants. This is because kallisto maps reads to the same region regardless of the allele: at heterozygous SNP loci, kallisto skips mapping the *k*-mer containing the SNP for efficiency, if the *k*-mer is located between tDBG junctions.

Ornaments was on average 26 times faster than WASP, and nearly as fast as kallisto (Fig. 4). Ornaments took 8.7 minutes per sample, similar to 8.4 minutes for kallisto, whereas WASP took 227.8 minutes, almost equally split between the initial read mapping and re-mapping for bias correction. The computation time of all methods was obtained on a single core of a machine with an Intel Xeon 2.5GHz processor and 32GB memory.

**Figure 4:**
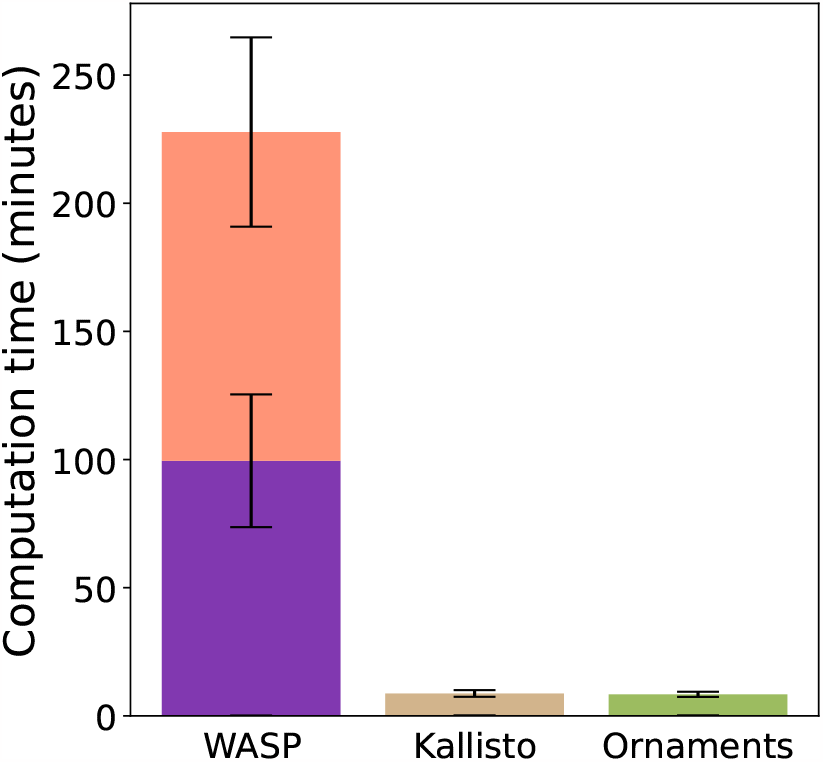
Computation time of Ornaments and other methods. Results averaged over 10 simulated samples are shown with error bars for one standard deviation. The computation time of WASP is split into the time taken to perform the initial read alignment using the STAR aligner (purple) and the time taken to run the WASP re-mapping pipeline (pink).

### Human lymphoblastoid cell-line RNA-seq reads

Using the lymphoblastoid cell-line reads and genome sequences for 165 children from the 1000 Genomes Project ^15^, we benchmarked Ornaments against WASP. We compared the allele-specific expression and allelic ratios from Ornaments with those from WASP, using gene-level summaries for Ornaments. Ornaments and WASP had consistently lower correlation at loci overlapping with ambiguously mapped reads than at the other heterozygous loci (Fig. 5), providing evidence for the superior ability of Ornaments to correct for ambiguous mapping bias.

**Figure 5:**
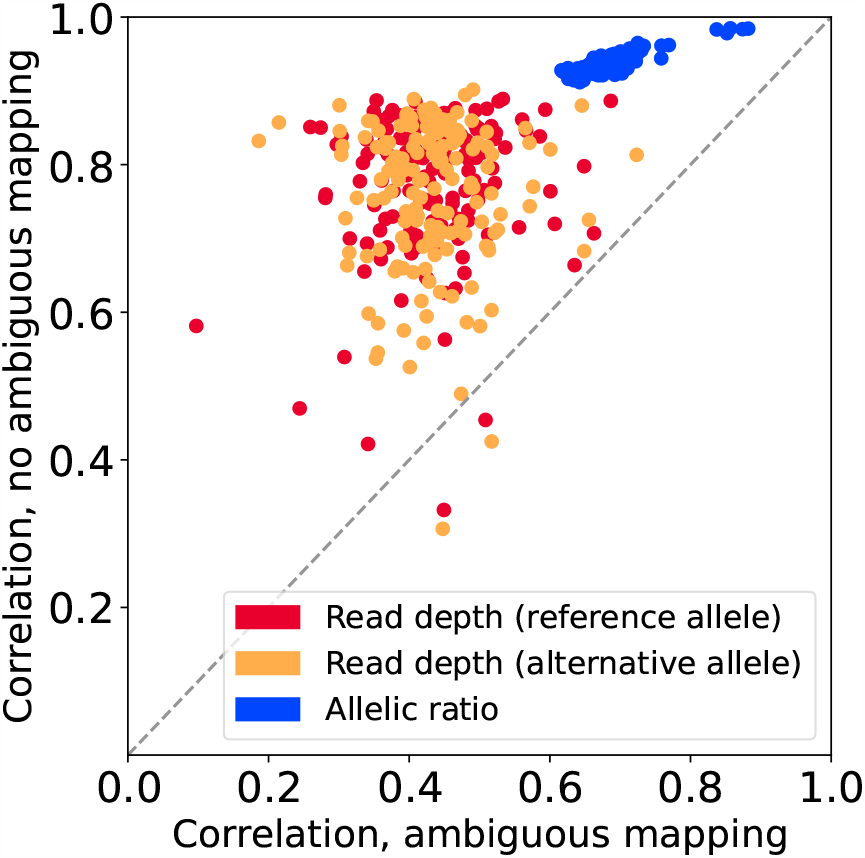
Comparison of Ornaments and WASP on lymphoblastoid cell-line RNA-seq reads. Correlation between Ornaments and WASP estimates across SNP loci with ambiguously mapped reads (*x*-axis) and without ambiguously mapped reads (*y*-axis). Each dot represents each of 165 samples.

In downstream analysis of differential expression of alleles with GeneiASE ^20^, Ornaments found the majority of the genes from WASP and a large number of additional genes. Ornaments found 4,374 genes with differentially expressed alleles in at least one constituent transcript of the gene in at least 10 samples, whereas WASP identified only 1,034 genes in at least 10 samples. Out of the 1,034 genes from WASP, 897 genes were also found by Ornaments. This suggests higher sensitivity of Ornaments in downstream analysis, as it finds allele-specific signals at transcript level with highly accurate bias correction.

To see if the expected read counts from Ornaments can reproduce known biological results, we examined an overlap between these genes with differentially expressed alleles and previously known imprinted genes, where one allele is exclusively expressed over the other. We compiled a set of 157 imprinted genes that are either known to undergo imprinting or found to be imprinted from an independent dataset. Our set included 141 genes in the GeneImprint database ^22^, 13 genes identified from analysis of lymphoblastoid cell-line RNA-seq data for 80 individuals with European ancestry ^23^ and for 63 unrelated individuals ^24^, and other known imprinted genes from the literature ^25,26,27^. Ornaments had 26 genes in the overlap, whereas WASP had a smaller overlap of 22 genes. Additionally, Ornaments detected a subset of transcripts of each gene exhibiting imprinting (Figs. 6, S6, and S7) and called differential expression in more samples for 17 out of the 21 imprinted genes found by both methods (Table 1). These results provide evidence that Ornaments with its probabilistic approach can accurately attribute the allele-specific signals to multiple transcripts of the given gene, providing advantages over WASP that captures gene-level signals.

**Table 1:**
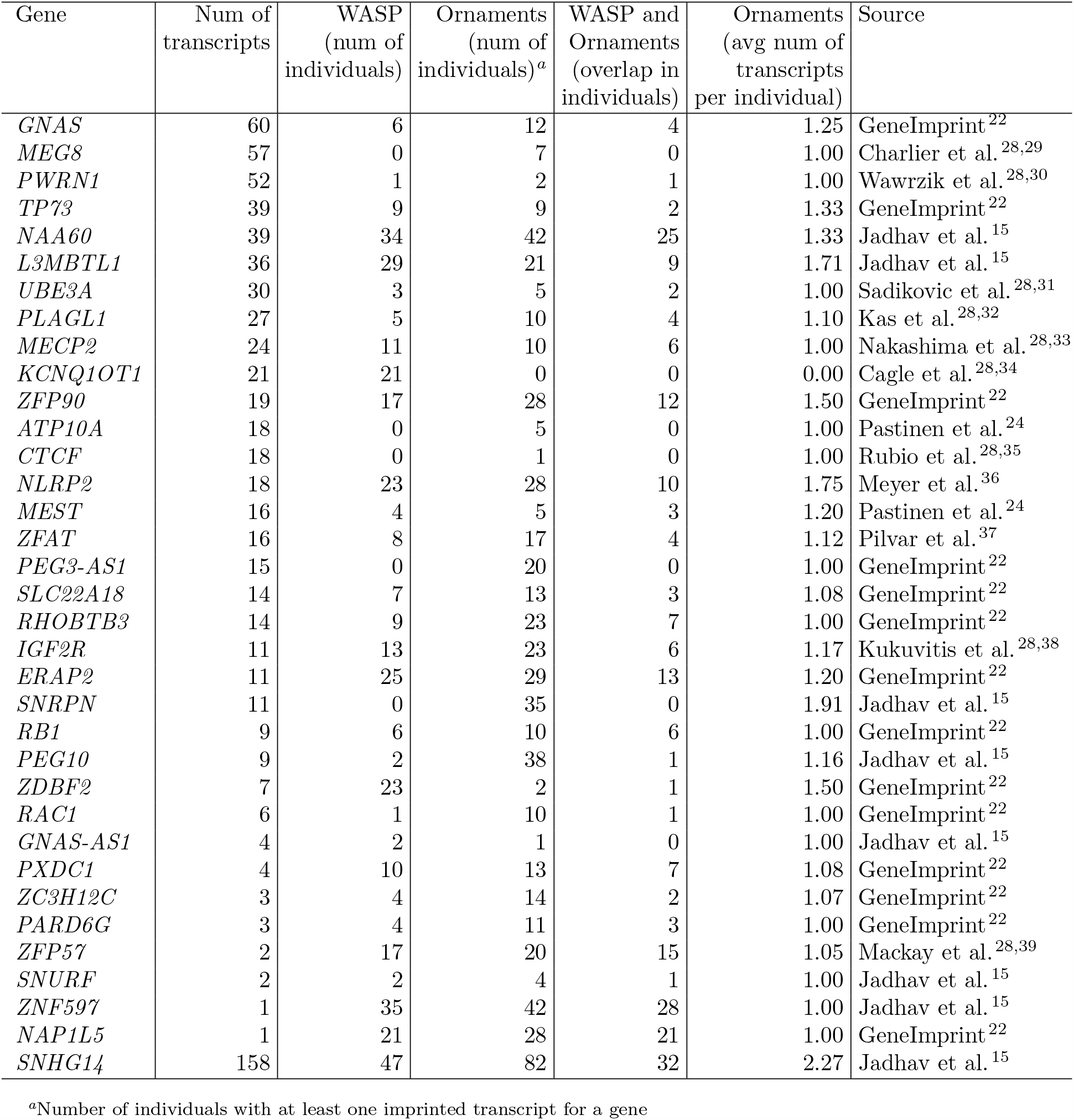
Known imprinted genes overlapping with allele-specifically expressed genes from WASP and transcripts from Ornaments in lymphoblastoid cell-lines.

**Figure 6:**
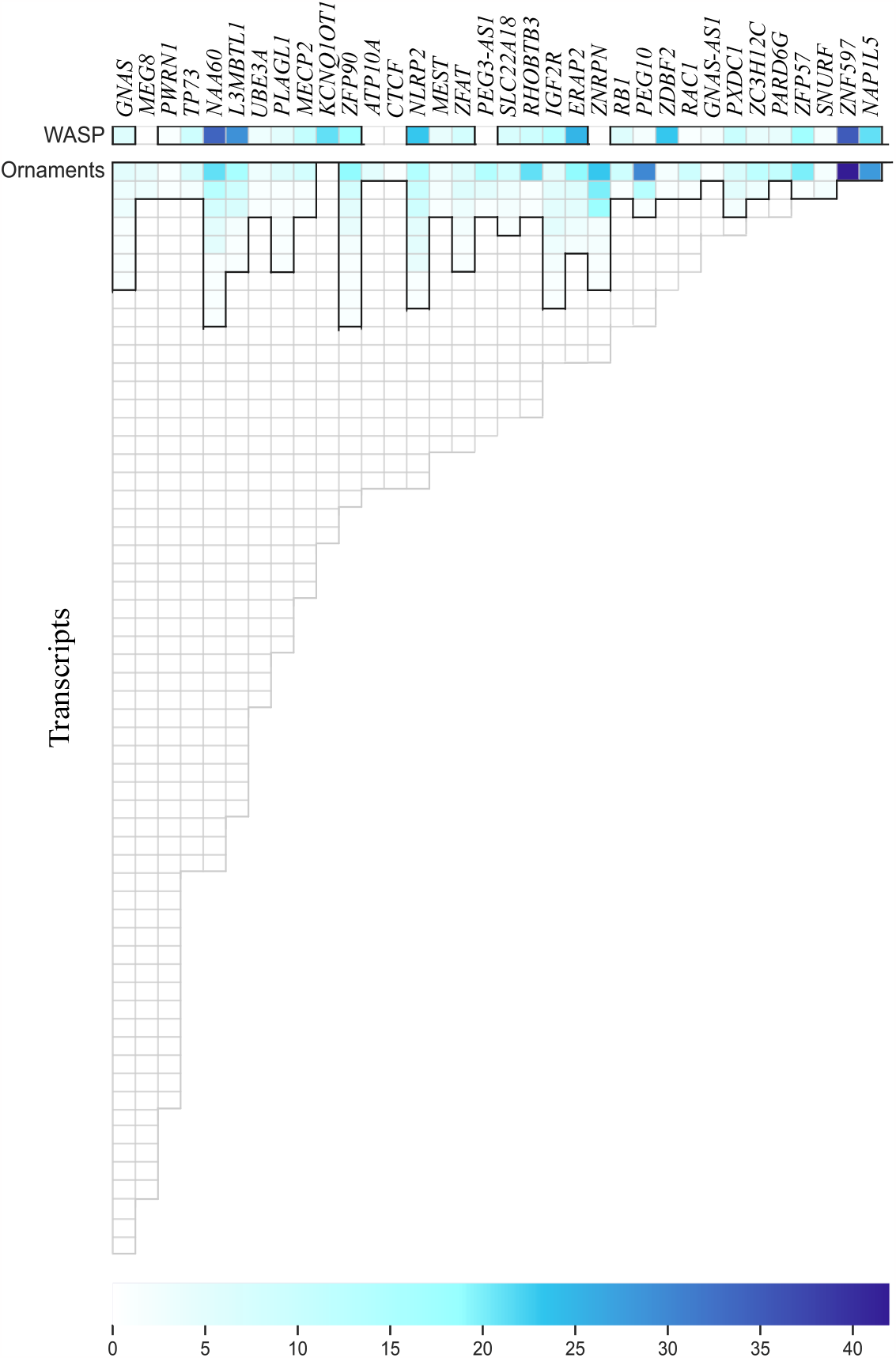
Known imprinted genes overlapping with allele-specifically expressed transcripts from Ornaments and genes from WASP in lymphoblastoid cell-line. Each cell shows the number of individuals found by each method for each transcript or gene. The cells with one or more individuals are enclosed in black lines. *SNHG14* with 158 transcripts, omitted in the figure, is a known imprinted gene that had allele-specifically expressed transcripts in 82 samples for Ornaments, but only in 47 samples for WASP.

## Discussion

We introduced Ornaments, a computational tool for accurate and efficient quantification of allele-specific expression at heterozygous loci from RNA-seq reads. Ornaments is a modification of kallisto that improves upon both kallisto and WASP. Ornaments improved the accuracy of WASP by probabilistically mapping allele-specific reads that WASP discards to correct for ambiguous mapping bias and by capturing allelespecific signals at transcript level rather than at gene level. Ornaments improved the accuracy of kallisto by accounting for variants during transcriptome quantification. In addition, Ornaments dramatically reduced the computation time of WASP by inheriting the speed of kallisto.

Future directions include further reducing computation time by processing multiple individuals in batches, when genotype data are available for a large number of individuals as in eQTL mapping. The resulting ornament tDBG would contain ornaments for all variants that are heterozygous in at least one sample in the batch. Then, when performing variant-aware pseudoalignment, we would make use of only those ornaments for variants that are consistent with the genotype for a given sample.

## Acknowledgments

This work was supported by NIH-1R21HG011116, NIH-1R21HG010948, and NSF-DBI2154089.

## Author contributions

A.A. and S.K. conceived the project, developed the method, designed the experiments, and wrote the paper. A.A. performed the experiments.

## Data and code availability

Contact the authors for the code.

## Supplemental Figures and Legends

**Figure S1:**
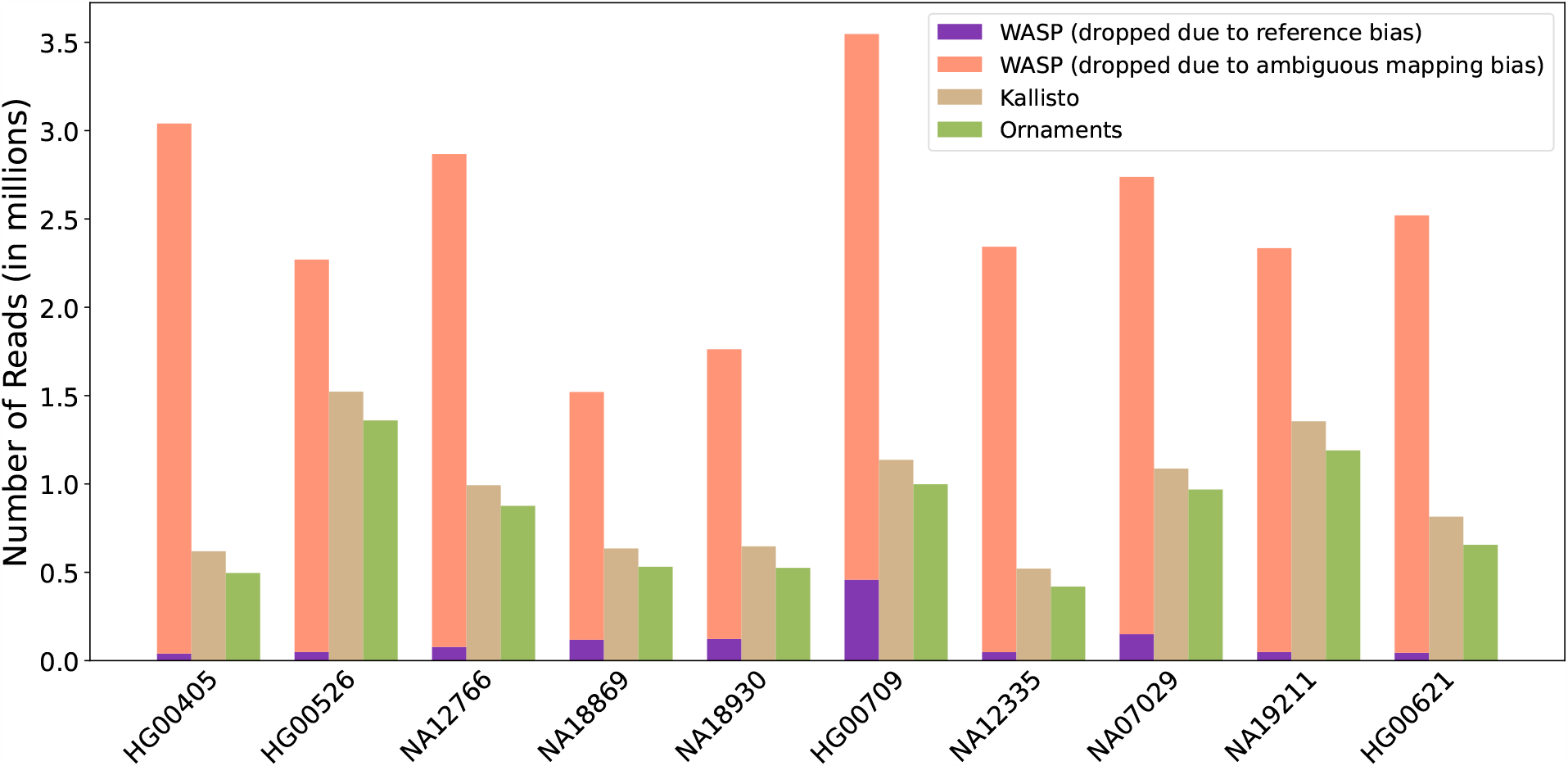
Number of reads dropped by WASP, kallisto, and Ornaments due to allele-specific read mapping bias in simulation study. The benchmark was performed on 60 million simulated reads for each sample. Reads that are shorter than length *k*=31 for *k*-mers in tDBG were dropped by all methods and were excluded.

**Figure S2:**
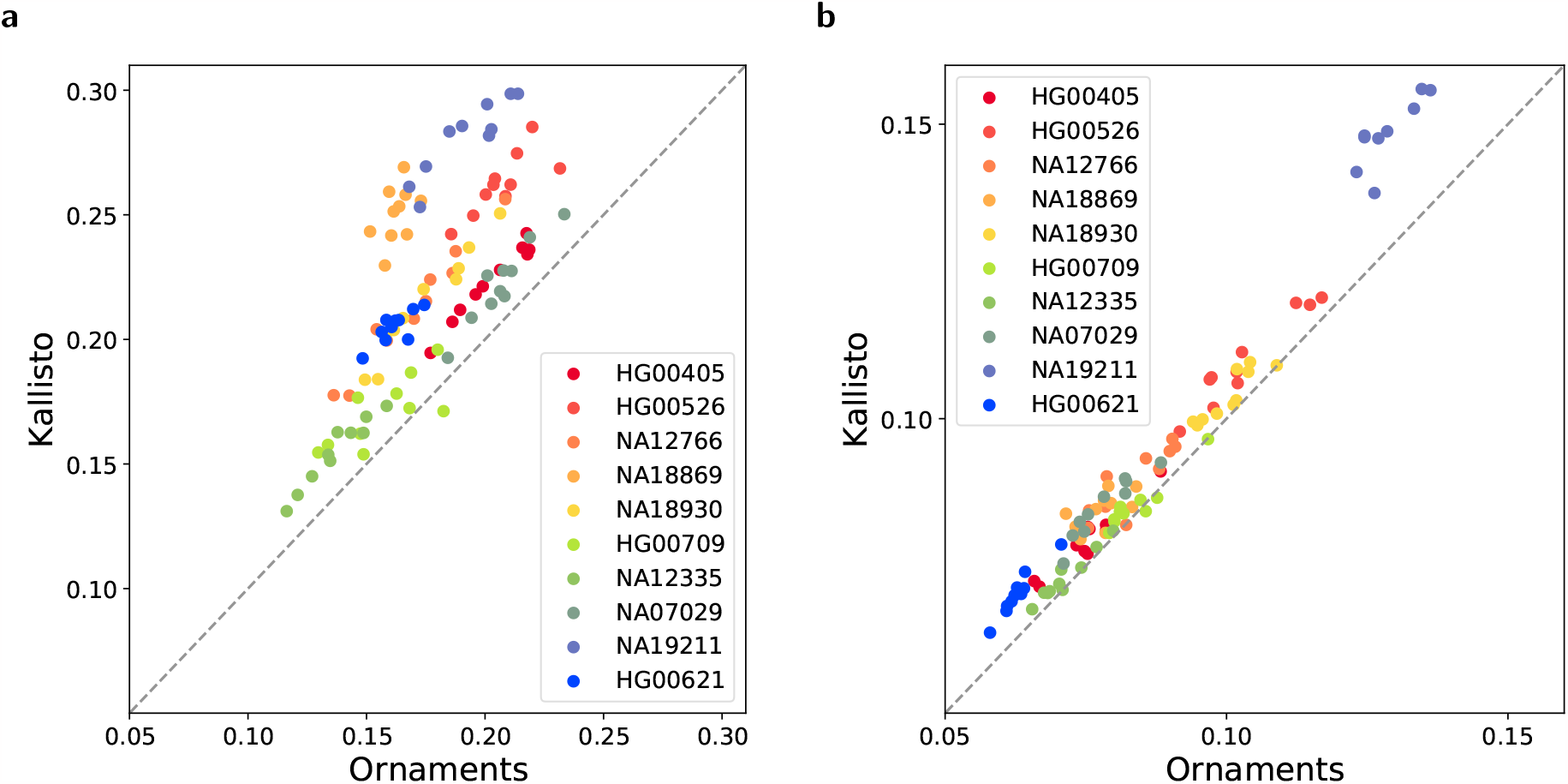
Comparison of Ornaments and kallisto on the accuracy of quantification of unexpressed transcripts containing ambiguously mapped reads in simulation study. The accuracy in mean absolute difference for (a) transcripts without SNPs and (b) transcripts with SNPs. Each dot represents the result from each of 10 repetitions of simulating reads from the genome of the given sample.

**Figure S3:**
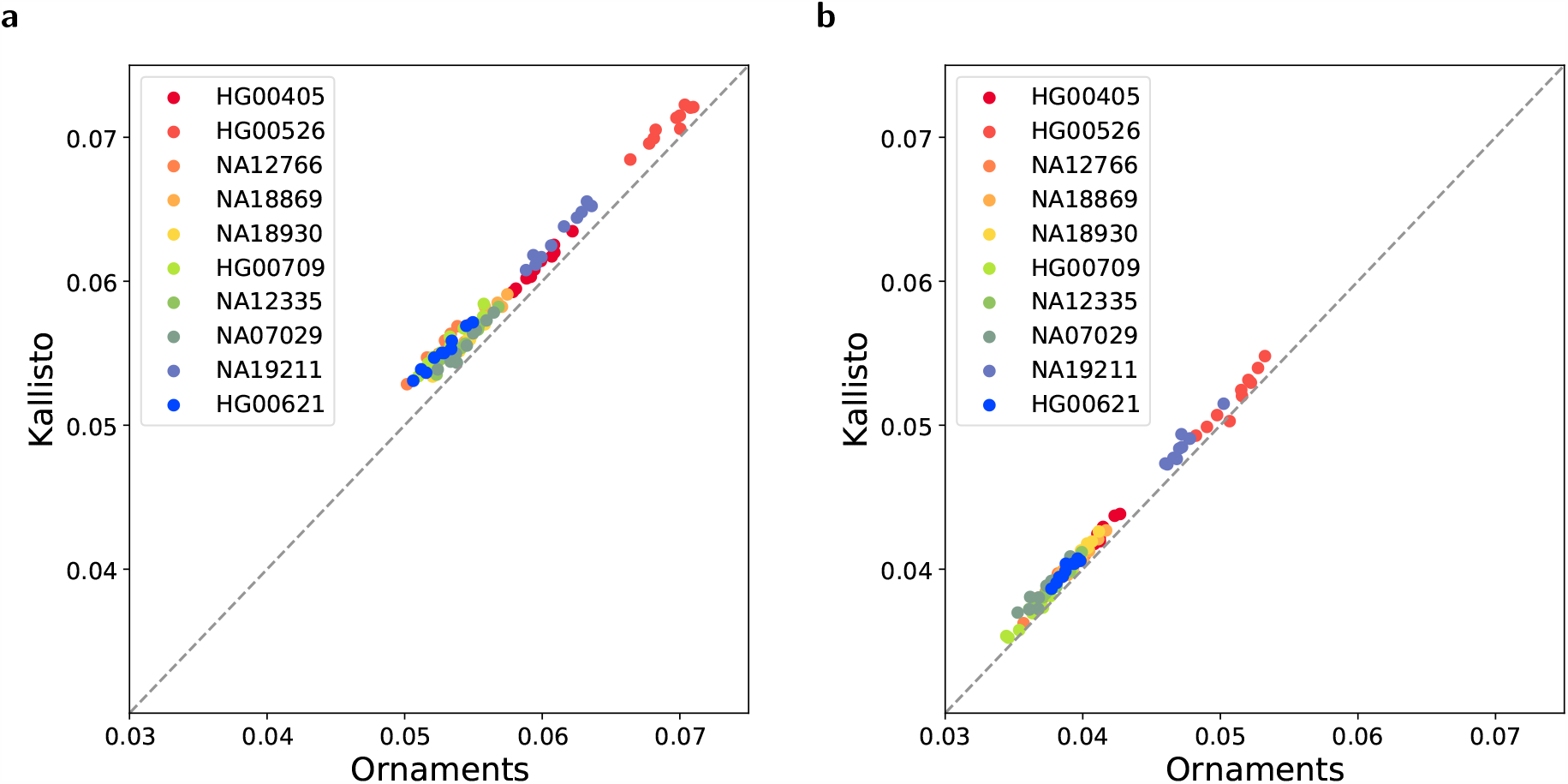
Comparison of Ornaments and kallisto on the accuracy of quantification of unexpressed transcripts containing no ambiguously mapped reads in simulation study. The accuracy in mean absolute difference for (a) transcripts without SNPs and (b) transcripts with SNPs. Each dot represents the result from each of 10 repetitions of simulating reads from the genome of the given sample.

**Figure S4:**
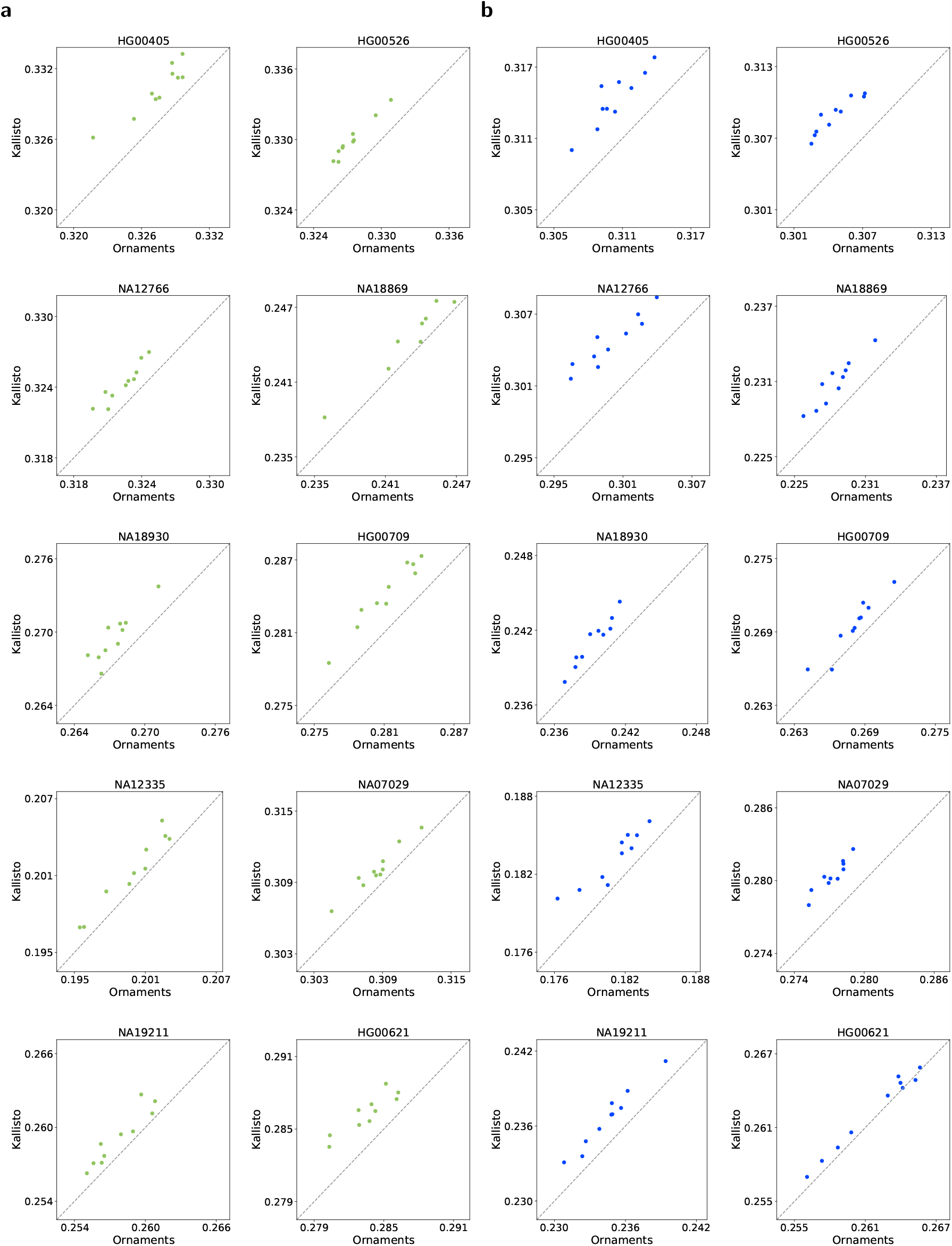
Comparison of Ornaments and kallisto on the accuracy of quantification of expressed transcripts containing ambiguously mapped reads in simulation study. The accuracy in mean absolute relative difference for (a) transcripts without SNPs and (b) transcripts with SNPs. Each dot represents the result from each of 10 repetitions of simulating reads from the genome of the given sample.

**Figure S5:**
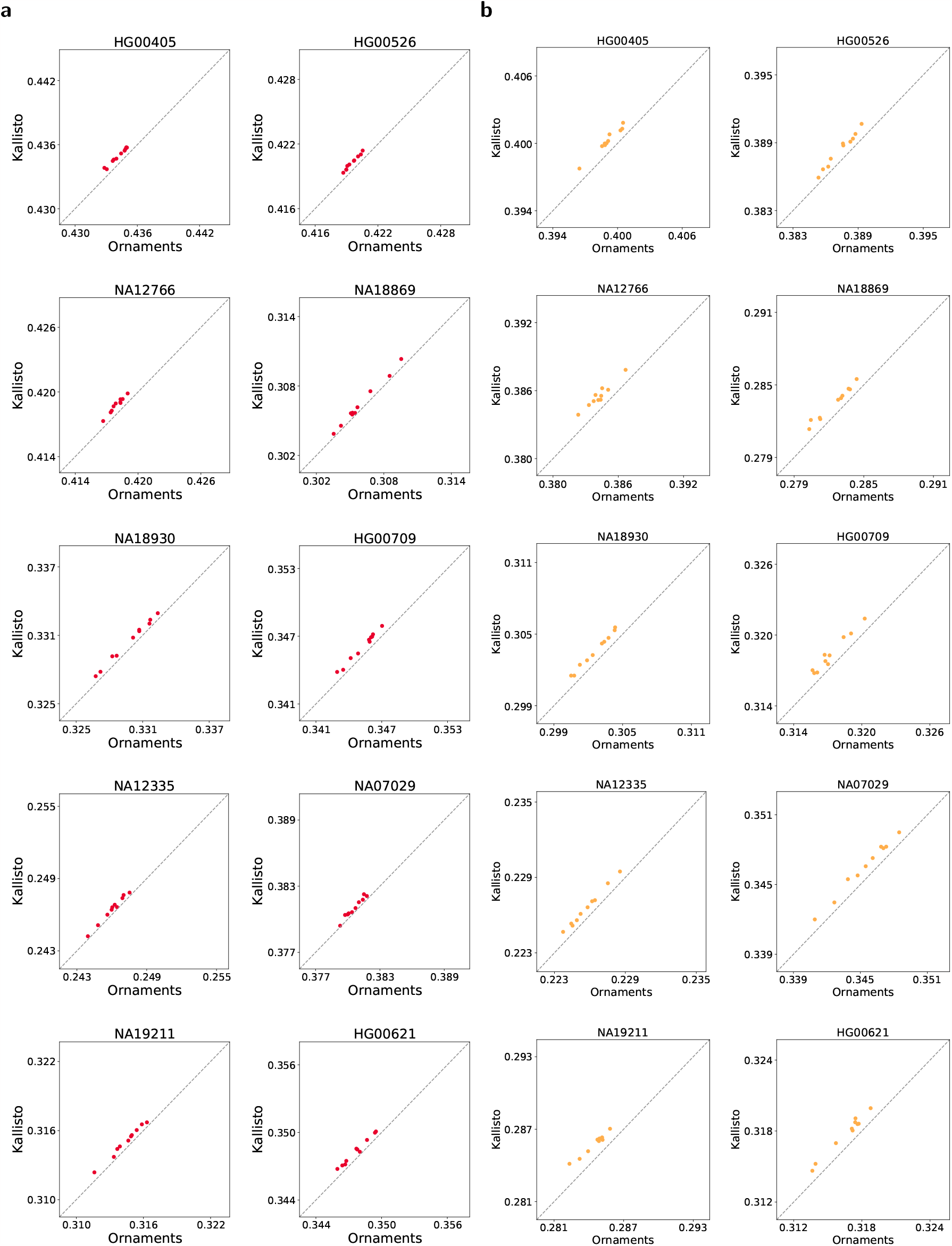
Comparison of Ornaments and kallisto on the accuracy of quantification of expressed transcripts containing no ambiguously mapped reads using simulated data. The accuracy in mean absolute relative difference for (a) transcripts without SNPs and (b) transcripts with SNPs. Each dot represents the result from each of 10 repetitions of simulating reads from the genome of the given sample.

**Figure S6:**
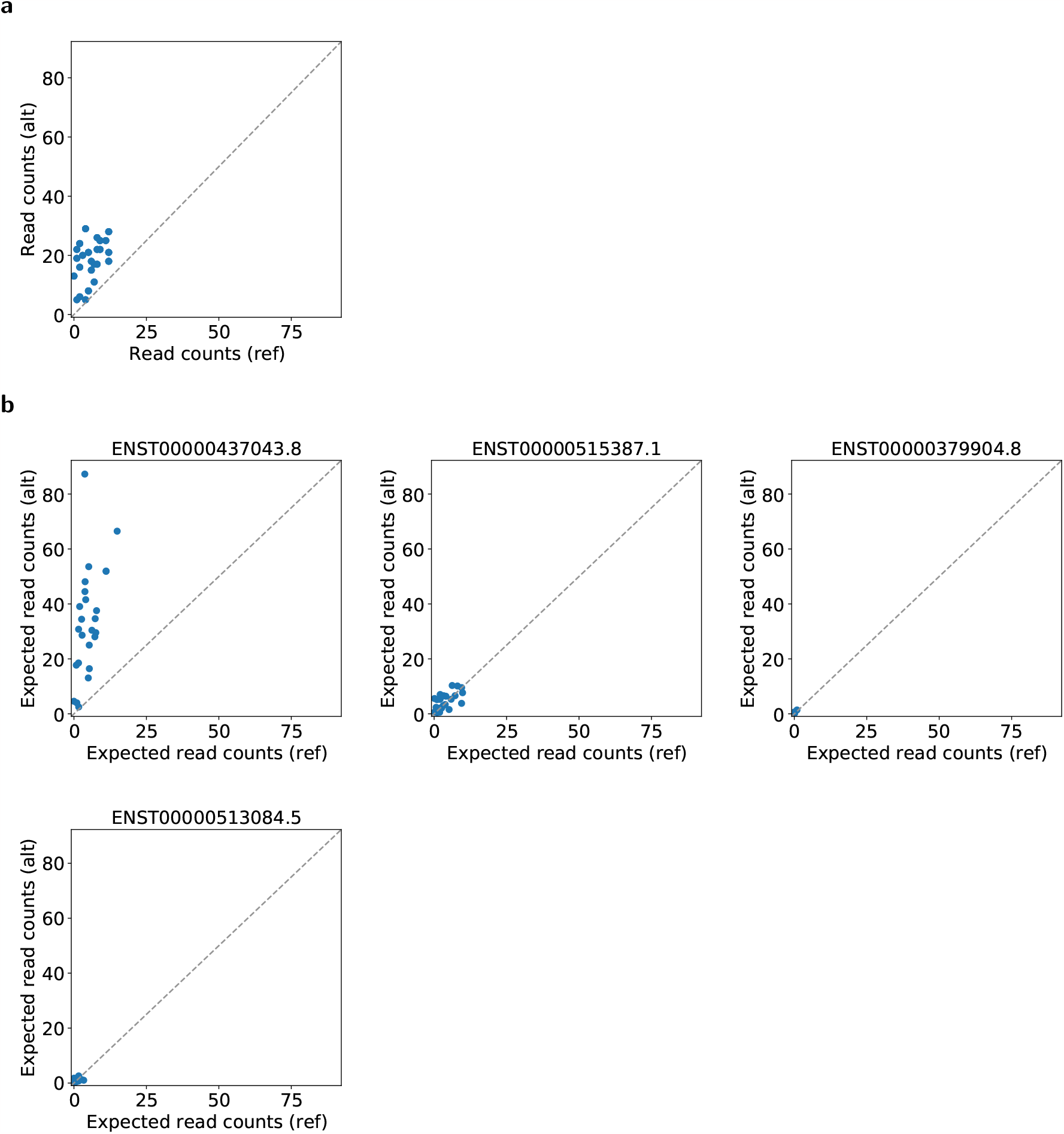
WASP vs Ornaments on allele-specific expression estimate at SNP rs1056893 in gene *ERAP2* using the lymphoblastoid cell-line reads. (a) Allele-specific read counts at gene level from WASP. (b) Expected allele-specific read counts at transcript level from Ornaments.

**Figure S7:**
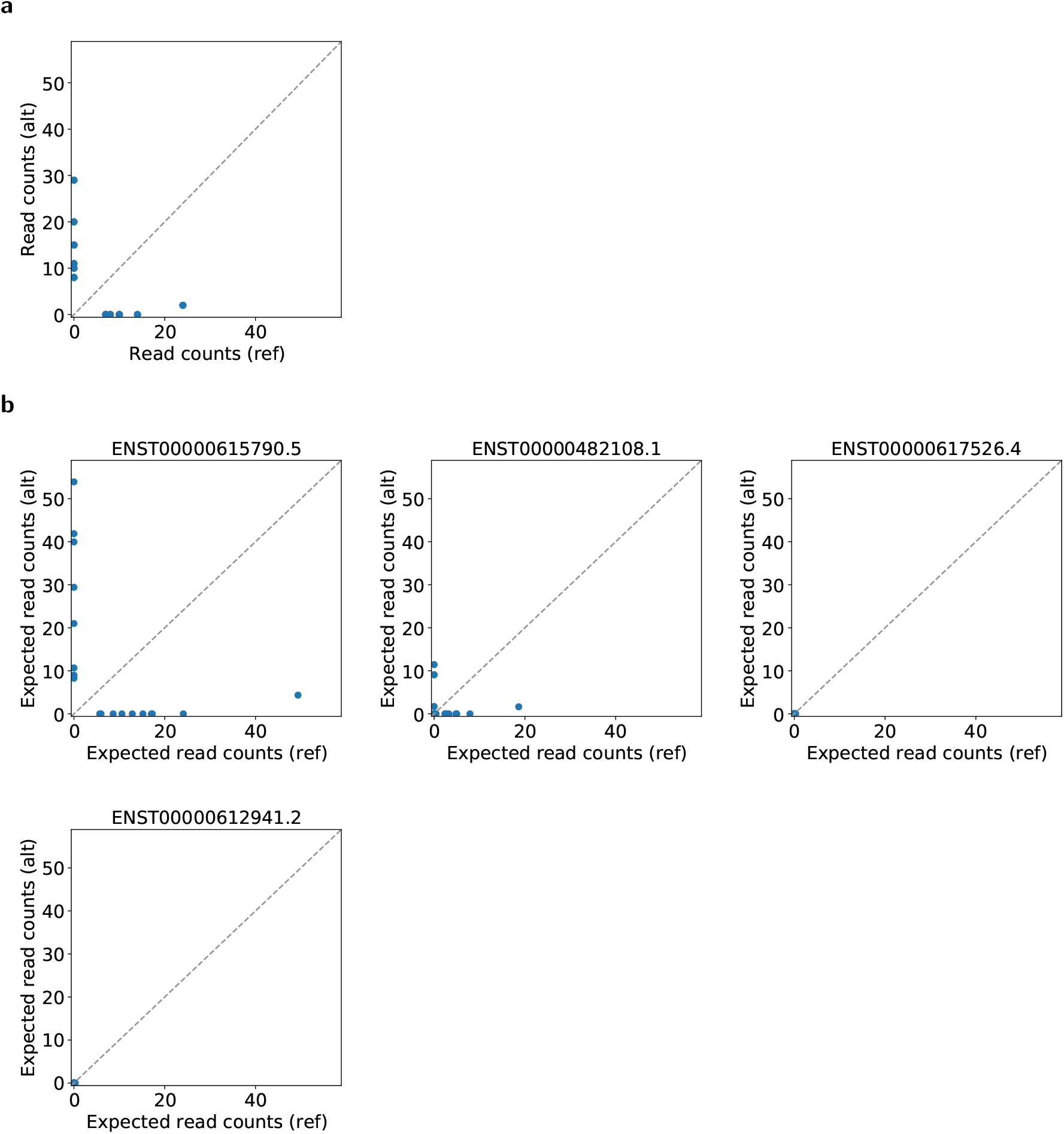
WASP vs Ornaments on allele-specific expression estimate at SNP rs13073 in gene *PEG10* using the lymphoblastoid cell-line reads. (a) Allele-specific read counts at gene level from WASP. (b) Expected allele-specific read counts at transcript level from Ornaments.

